# Metabolic interactions of a minimal bacterial consortium drive robust nitritation at acidic pH

**DOI:** 10.1101/2023.10.29.564480

**Authors:** Gaofeng Ni, Zicheng Su, Yu Wang, Zhiyao Wang, Mengxiong Wu, Zhengshuang Hua, Shihu Hu, Zhiguo Yuan, Jianhua Guo, Chris Greening, Min Zheng

## Abstract

Microbial communities efficiently mediate aerobic ammonia oxidation even at acidic pH. However, little is known about the adaptations and interactions that allow these communities to withstand challenges such as acidic stress, reactive nitrogen species, and resource deprivation under such conditions. Here we combined metagenomic analysis and biogeochemical measurements to infer the composition, metabolic interactions, and stress adaptation mechanisms of microbial consortia in three acidic nitritation bioreactors, operating at pH 5 to 2. This resulted in the recovery of 70 high-quality and mostly novel metagenome-assembled genomes (MAGs). The dominant ammonia oxidiser across all three bioreactors was a novel proteobacterium, herein named Candidatus (*Ca*.) Nitrosoglobus kelleri, that we enriched to a relative abundance of 55%. Also present were several heterotrophic bacteria that are predicted to engage in metabolically cross-feeding with the *Nitrosoglobus*. Particularly abundant were mycobacteria, including the novel actinobacterium *Ca*. Mycobacterium cookii, which are predicted to use organic carbon, hydrogen, carbon monoxide, sulfide and possibly nitrite as energy sources to drive aerobic respiration and denitrification. Remarkably, we observed efficient conversion of ammonia to nitrate even at pH 2, by a minimalistic community comprising the *Nitrosoglobus* and *Mycobacterium* as its only core members. Genomic analysis suggests these bacteria each use multiple strategies to maintain intracellular pH homeostasis, detoxify reactive nitrogen species, and scavenge nutrients at this pH. Altogether, these findings reveal that minimal communities can drive a key biogeochemical process even at acidic pH, and have implications for understanding nitrogen cycling and enhancing wastewater treatment.

## Introduction

Ammonia oxidizing bacteria (AOB) and ammonia oxidizing archaea (AOA) rely on a suite of physiological adaptations to maintain pH homeostasis, acquire sources of energy and carbon, and detoxify reactive nitrogen species to mediate nitrification at acidic pH^1^. Despite the multifacetedness and complexity of the challenges to carry out acidic nitrification, diverse microorganisms have been reported to mediate this process, sparking wide research interest^1–6^. However, our current understanding of this process has primarily been informed by genomic inferences from a limited number of characterised nitrifiers. The acidophilic AOA *Candidatus* (*Ca*.) Nitrosotalea devaniterrae uses high affinity substrate acquisition systems coupled with mechanisms for ion transport, proton consumption and membrane adaptation to grow and oxidize ammonia at pH 4 – 5.5^4^. The acid-tolerant AOB *Ca*. Nitrosoglobus terrae TAO100 (herein referred to as TAO100) employs cation transporters, urease, and glycosyltransferase to confer acid-tolerance^5^. Its cells aggregate into biofilms, mediated by the synthesis of exopolysaccharide to survive at pH as low as 2.0^5^. Similarly, another acid-tolerant AOB, the *Ca.* Nitrosacidococcus tergens sp. RJ19 (herein referred to as RJ19), possesses 12 cation transporters^6^, which creates a reverse membrane potential to repel proton intrusion and regulate pH homoestasis^7^. Among them, one Na^+^/H^+^ antiporter shares homology with antiporters responsible for regulating pH homeostasis in *E. coli* K12,^8,9^. Also, the presence of two cytoplasmic carbonic anhydrases in the genome of TAO100 mediates proton scavenging and pH homeostasis^5^, as seen in the obligate acidophilic ammonia oxidizing archaea *Ca.* Nitrosotalea devaniterrae^2,4^. Finally, reactive nitrogen species such as nitric oxide (NO) or nitrite (NO ^-^) produced through ammonia oxidation are believed to be mediated by NO reductase in RJ19^6^. However, since nitrifiers rarely function by themselves, the metabolic interactions between the chemoautotrophic nitrifiers and the neighboring heterotrophs that allows them to adapt and colonise these ecosystems is a crucial area that needs to be addressed. This will provide a comprehensive and in-depth picture of this significant nitrogen transformation process in natural, agricultural, and engineering settings.

Recently, acid-tolerant AOB from *Ca*. Nitrosoglobus have been suggested to drive acidic nitritation reactor systems, which are wastewater treatment processes operating under an acidic pH (e.g., 4.5–5.0) ^10,11^. This novel process enables nitrite accumulation under acidic conditions, which generates protonated form of nitrite (HNO_2_), a strong biocidal compound termed “Free Nitrous Acid (FNA)” that severely suppresses the growth of nitrite oxidizing bacteria (NOB).

NOB suppression enables the coupling of acidic nitritation with anaerobic ammonium oxidation (anammox)^10,12^, which reduces energy consumption from aeration by 60% while completely eliminating the need to dose organic carbon to sustain the functioning of heterotrophic denitrifiers in the activated sludge treatment process^13^. The activated sludge process is the world’s largest biotechnology due to the fact that it is applied globally in wastewater treatment plants (WWTPs) and its treatment volume equates to one-seventh of global river flow^14^. However, WWTPs demand 3 – 5% of national energy budgets in developing and developed countries^15,16^, making it an energy-intensity industry. Successful adoption of acidic nitritation holds tremendous potential to reduce the environmental footprint from WWTPs. However, the realisation of this potential hinges on a comprehensive and in-depth understanding of its keystone microorganisms, a knowledge area currently insufficiently explored.

Previously, amplicon sequencing variants (ASVs) obtained from 16S rRNA gene amplicon analysis confirmed that the key nitrifier driving acidic (pH 2-5) nitritation in laboratory-scale bioreactor systems are *Ca*. Nitrosoglobus spp.^10,17,18^. Upon obtaining an enriched culture at 55% relative abundance, stoichiometric and kinetic characterization have shown that the enriched *Ca*. Nitrosoglobus spp. bears remarkable tolerance to pH as low as 2 and FNA at 2 - 8 ppm, which are the most adverse environmental conditions known to nitrifiers in this regard^17^. These traits define the *Ca*. Nitrosoglobus as an adversity-strategist^19^ that thrive at acidic activated sludge ecosystems. Previous work has shed light on the physiological, ammonia oxidation kinetics and genomic characteristics of *Ca*. Nitrosoglobus terrae TAO100, which was enriched from tea fields^5^, and *Ca*. Nitrosacidococcus tergens RJ19, which was discovered from an air scrubber treating exhaust air from a pig stable^6^. However, within acidic nitritation reactor systems, genome-resolved understanding of *Ca*. Nitrosoglobus populations as well as its interaction with heterotrophic microorganisms is absent. Furthermore, isolation of acid-tolerant AOB has been challenging, with only limited enrichment cultures available^5,17^. This leads us to hypothesize that a metabolic partnership between acid-tolerant AOB and its closely associated heterotrophic partners is crucial for its growth, a premise that is currently untested. To this end, we have coupled bioreactor operation with high throughput metagenomic sequencing analysis in three lab-scale acidic nitritation bioreactors. Our study has generated an unambiguous genome-resolved understanding of metabolic partnership that facilitates acidic nitritation.

## Methodology

### Bioreactor operation

To achieve stable nitritation at acidic pH and obtain acid-tolerant AOB, a membrane bioreactor (designated R1, working volume 2L) was operated, which contains a microfiltration membrane module to retain biomass^10^. It was continuously fed with municipal wastewater containing 92 mg N/L ammonia and was operated at pH between 4.5 and 5.0^10^. To enrich the acidic tolerant AOB obtained from R1, membrane bioreactor R2 (10L) was inoculated using biomass from R1 and was fed with synthetic wastewater containing ammonia between 150 – 250 mg N/L and no organic carbon^17^. A programmable logic controller (PLC) was implemented to control R2 pH at 5.0^17^. To explore lower pH limit for ammonia oxidation, a sequencing batch reactor R3 (2L) was operated using biomass from R2 and was fed with anaerobic digestion liquor (AD liquor), which contains much higher level of ammonia (881 mg N/L), and a pH of 2.0 was achieved through the intrinsic ammonia oxidation without pH controlling^18^. The concentration of nitrogen species from reactor liquids was measured using a Flow Injection Analyzer (Lachat QuickChem8000, Milwaukee)^10,17,18^.

### DNA extraction

To extract DNA, 0.5 mL of biomass (suspended sludge) was sampled from the reactors during stable acidic nitration. DNA was extracted using FastDNA® spin kit (MP Biomedicals, Santa Ana, CA) following the manufacturer’s protocol. DNA concentration was assessed using a NanoDrop spectrophotometer (Thermo Fisher Scientific, United States).

### Metagenomic sequencing and the recovery of MAGs

Metagenomic sequencing was performed at the Australian Centre for Ecogenomics at the University of Queensland, on a NextSeq 500 platform (Illumina, U.S.), resulting in 96.0, 112.6, and 110.7 million read pairs (2 x 150 bp) from the R1, R2, and R3, respectively. Read quality control, assembly, binning, bin refinement and reassemble were carried out in a metagenomic wrapper tool MetaWRAP v1.2.2^20^.

Raw metagenomic reads were trimmed based on adaptor content and PHRED scores (minimum 20) in Trim-galore v0.5.0^21^. Individual assembly from each sample was carried out using metaSPAdes v3.13.1 with kmers of 21, 33, 55, 77, 99, and 111^21^. Bins from three binning strategies, i.e., MaxBin2 v2.2.4^22^, metaBAT2 v2.12.1^23^, and CONCOCT v1.0.12^24^ were consolidated into one superior set of MAGs that has at least 70% completeness and a maximum of 10% contamination. Afterwards, MetaWRAP “reassemble_bins” module was utilized to improve N50, completeness and to reduce contamination of the final bin set, which yielded 70 high-quality metagenomic assembled genomes (MAGs). CheckM2 “predict” workflow v1.0^25^ was used to verify genome completeness and contamination. The relative abundance of each MAG in its respective reactor was estimated using CoverM v0.6.1^25^.

### Taxonomic annotation, functional profiling, and phylogenetic analysis

Taxonomic classification of the MAGs was performed using GTDB-tk v2.1.1^26,27^ against the Genome Taxonomy Database (GTDB, R207). Subsequently, open reading frames (ORFs) from each MAG were called and a primary functional annotation was carried out using Prokka (v1.4.0)^28^. Metabolic profiling for nitrogen metabolism, sulfur metabolism, and respiration in both short reads and MAGs was carried out using Diamond v2.0.14^29^ to search against the Greening Lab metabolic marker gene database^30–34^. The presence of marker genes associated with microbial carbon, nitrogen, and sulfur metabolism, trace gas oxidation, and respiration was identified for MAGs affiliated with *Ca*. Nitrosoglobus and *Mycobacterium* using manually curated threshold for homology-based mapping^35^. The presence and absence of these genes were also summarized for the families Chitinophagaceae, Sphingomonadaceae, Rhodanobacteraceae and Burkholderiaceae. Annotations of carbohydrate active enzymes (CAZymes) in each MAG was achieved based on search against the CAZyme HMM database (dbCAN HMMdb r8.0) following the dbCAN2 CAZyme annotation workflow^36^. Heatmap of the count of CAZyme in each MAG according to each category i.e., glycoside hydrolases (GHs), glycosyl transferases (GTs), polysaccharide lyases (PLs), carbohydrate esterases (CEs), carbohydrate binding modules (CBMs), and auxiliary activities (AAs)^37^ was generated using the python packages seaborn^38^ and matplotlib^39^.

Phylogenomic tree of the 70 MAGs recovered in this study is generated using Phylophlan 3^40,41^, which incorporates multiple sequence alignment with MAFFT^42^, alignment trimming with Trimal v1.2rev59 (“-gappyout” option)^43^, and tree inference in IQ-TREE2 v1.6.12 (“-alrt 1000-B 1000 -m TEST” option for automatic model testing and bootstrapping with 1,000 iterations)^44^. The ETE3^45^ toolkit and ITOL^46^ for tree rendering. To closely examine *Ca*. Nitrosoglobus and *Mycobacterium* populations, the phylogenomic tree file generated from the “classify_wk” in GTDB-tk during taxonomic assignment was visualized in Seaview^47^, and GTDB species representatives that are close to *Ca*. Nitrosoglobus and *Mycobacterium* MAGs of this study were manually selected and included in the phylogenomic analysis using Phylophlan3^40^. Furthermore, *Methylophaga lonarensis* MPL (GCF_000349205.1) and *Corynebacterium diphtheriae* (GCA_001457455.1) were used as outgroups during phylogenomic reconstruction for Nitrosococcaceae and *Mycobacterium*, respectively, because they were the closest outgroup lineages. Similarly, a phylogenetic tree of the ammonia monooxygenase subunit A (AmoA) was generated using MAFFT^42^ for multiple sequence alignment, followed by trimming in Trimal^43^, and IQ-TREE2^44^ using the same parameters stated above. Sequences of particulate methane monooxygenase subunit A (PmoA) from Methylococcales (phylogenetically parallel Nitrosococcales within Gammaproteobacteria) were used as an outgroup. For comparative analysis within the family Nitrosococcaceae, Anvi’o^48^ was used to extract gene clustering, and gene clusters exclusive to acid-tolerant AOB were annotated against KEGG^49^, Pfam^50^ and COG databases^51^.

## Results and discussion

### Nitrification performance under acidic conditions

All three reactors carried out stable nitrification for a duration of 100 days, with average ammonium removal efficiencies of 83.6%, 57.0%, and 56.8% for R1, R2, and R3, respectively. This confirmed effective ammonium oxidation through nitrification at pH 2 to 5. Nitrite (NO_2_ ^-^) was the primary nitrification end product in R1 and R2, confirming that stable nitritation was achieved. In R3, nitrate (NO ^-^) was the predominant nitrification end product. This is likely attributable to the enhanced chemical oxidation of nitrite under extremely low pH conditions^6,52^.

### A novel and ultra-simplistic microbial community underpins acidic nitritation

Three MAGs affiliated with Ca. Nitrosoglobus were recovered from R1, R2 and R3 respectively (R1-bin45, R2-bin10 and R3-bin1), and were placed within Nitrosococcaceae based on in-depth phylogenomic analysis (**Fig**. 2, S1A). Furthermore, phylogenetic analysis of the ammonia monooxygenase subunit A (AmoA) extracted from these three genomes confirmed this placement (**Fig**. S1B). Also, comparative genomics analysis showed that the genomes of *Ca. Nitrosoglobus kelleri* shares between 91.7% to 91.8% ANI with *Ca.* Nitrosoglobus terrae TAO100; they form a distinct cluster from TAO100 and other representative species within Nitrosococcaceae, confirming that *Ca. Nitrosoglobus kelleri* is a novel species of *Ca*. Nitrosoglobus. The ANI among the three *Ca. Nitrosoglobus kelleri* genomes were between 98.4% and 99.7%, suggesting strain-level diversity (**Fig**. S1C). Their genome sizes are 2.0 MB, and the GC content is 42.1%, which is similar to that of TAO100 ^5^. These three genomes represent the third acid tolerant AOB species to date.

**Fig. 1.**
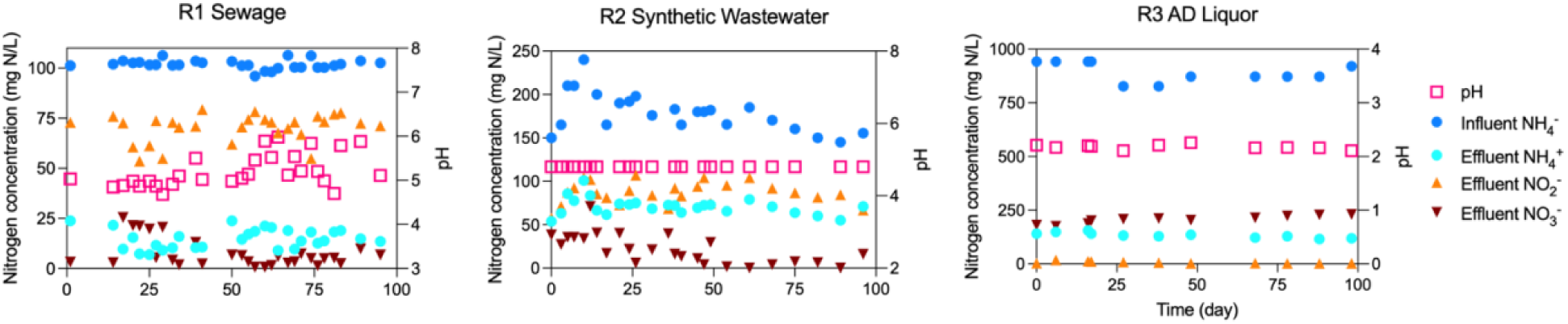
The performance of the three reactors regarding concentration of nitrogen species in the reactors expressed in mg/L (left Y axis) and pH (right Y axis).

**Fig. 2.**
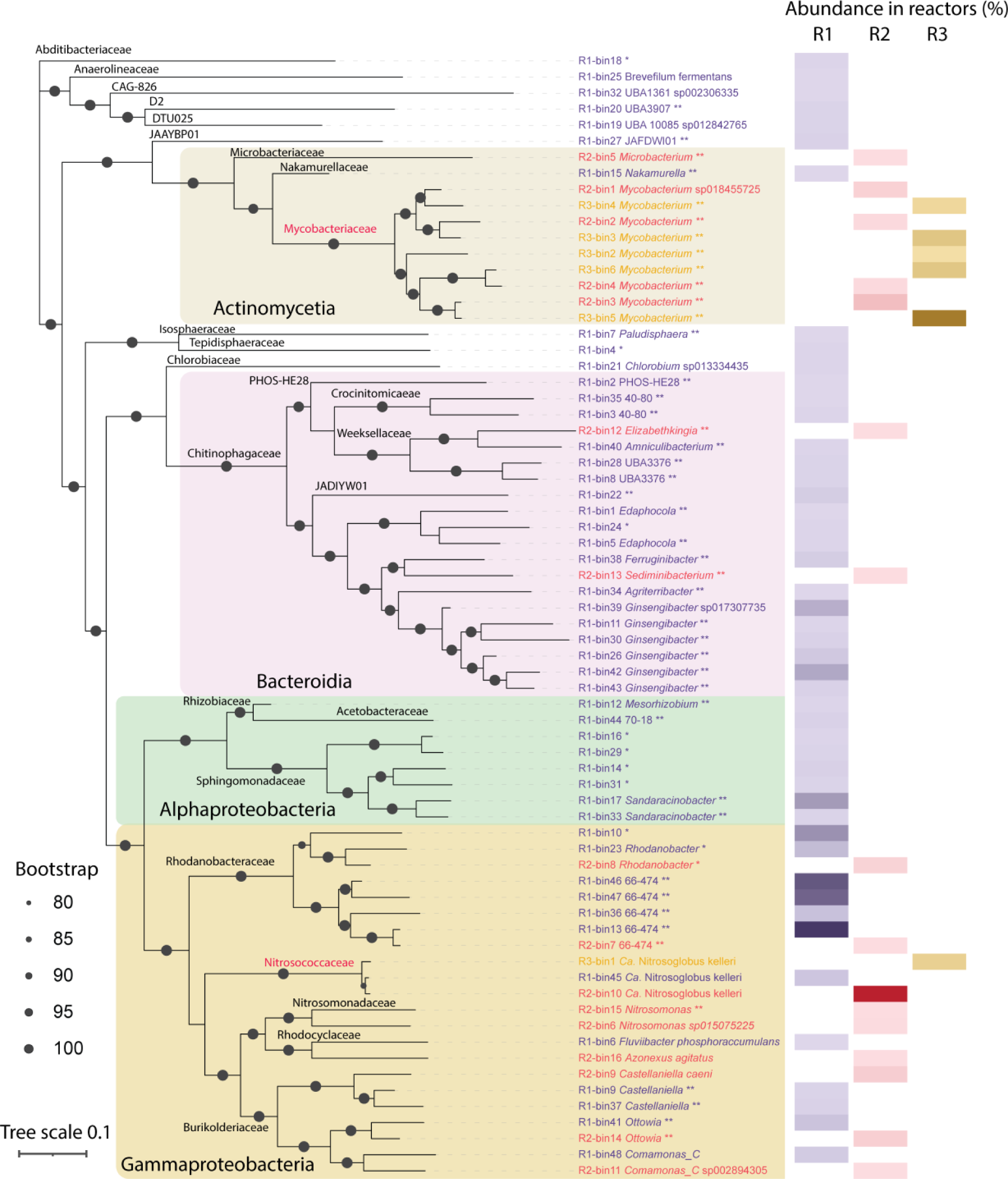
**Left**: a phylogenomic tree inferred from 400 concatenated universal proteins alignment showing the phylogenetic diversity of the 70 MAGs with class-level taxonomy assignment in coloured boxes, family-level taxonomy near nodes, and genus and species assignments on branches. The color of leaves indicates the reactor: R1 (purple), R2 (red), and R3 (brown). *: organism with unclassified genus; ** organisms with unclassified species. **Right**: a heatmap of relative abundance of MAGs in reactors, using the same color scheme to differentiate reactors.

Diverse *Mycobacterium* populations represented by nine MAGs were recovered from R2 and R3 (**Fig**. S2). Specifically, R2-bin1 and R3-bin4 were closely affiliated with *Mycolicibacter* (synonymous to *Mycobacterium*) *acidiphilum* M1^T^, which was isolated from a bioreactor treating high-ammonia wastewater at pH 2^53^. These MAGs are phylogenetically close to, yet distinct from the *Mycobacterium terrae* clade^54^ (**Fig**. S2). They reveal the under-appreciated diversity of *Mycobacterium* that are able to persist at low pH and high nitrosative stress.

In terms of relative abundance, diverse Rhodanobacteraceae from Gammaproteobacteria constitute 50.4% of the microbial community, while the sole AOB - *Ca*. Nitrosoglobus kelleri represented 1.4% of the community in R1. This suggests that *Ca*. Nitrosoglobus is highly active despite being low in abundance in R1. Reactor R2 was set out to enrich *Ca*. Nitrosoglobus kelleri, which reached a high abundance 58.4%. It coexisted with *Mycobacterium* (16.3%) and other members of Gammaproteobacteria (12.5%). Most strikingly, at pH 2, a condition characterized by extremely high acidity and nitrosative stress^17^ - an ultra-simplistic community with two species being *Mycobacterium* (57.0%) and *Ca*. Nitrosoglobus kelleri (7.2%) was enriched (**Fig**. 2). This strongly suggests a mutualistic partnership between these two species, enabling them to thrive under extremely low pH and high nitrosative stress conditions, while carrying out nitrification and aerobic respiration as a community.

Additionally, MAGs derived from the reactor biomass represent eight phyla in total. On the class level, Gammaproteobacteria were the most prevalent, followed by Actinomycetia and Bacteroidia according to the GTDB classification^55^ (**Fig**. 2). This observation deviates from global surveys of activated sludge ecosystems, where Betaproteobacteria overwhelmingly dominate over other taxonomic lineages^56^. Among the 70 high-quality (**Fig**. 3A) MAGs recovered in this study, 59 are without close relatives of known microorganisms (defined as having ANI radius of 80% within the GTDB database r207). Two MAGs representing *Nitrosomonas* were recovered from R2, they accounted for less than 1% of the community, it is likely that *Nitrosomonas* can adapt to acidic conditions^57,58^, yet their contribution to acidic nitritation is limited considering that they were dominated by *Ca*. Nitrosoglobus kelleri. Overall, our findings suggest that taxonomically novel microbial consortia underpin acidic nitritation systems.

**Fig. 3.**
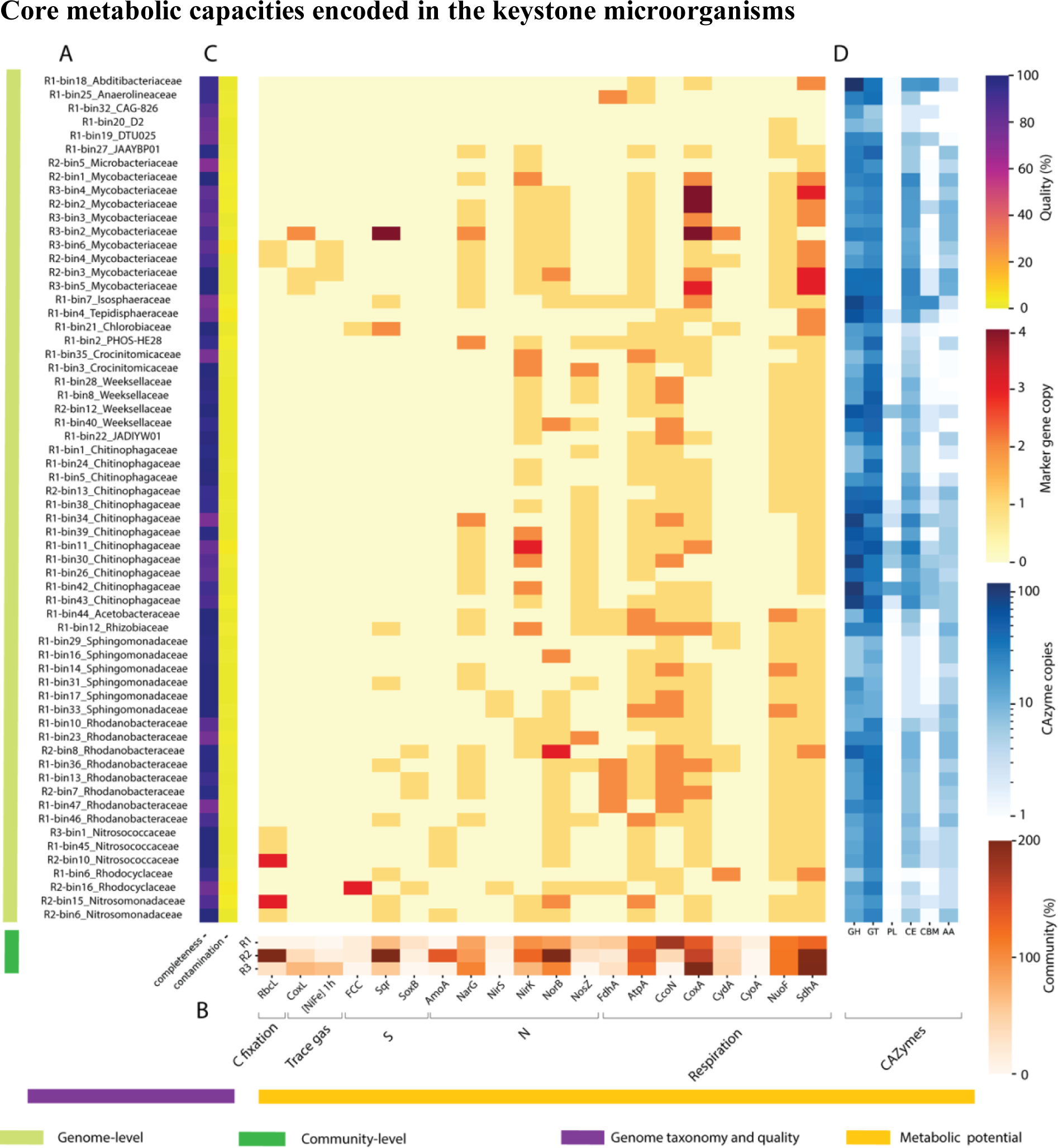
**A**: Heatmap showing genome completeness and contamination levels estimated using CheckM2. **B**: Heatmap showing the relative abundance of metabolic marker genes encoded in each of the three reactors, expressed as average gene copy per organism calculated relative to the abundance of 14 universal single-copy ribosomal genes. Metabolic categories include carbon fixation, trace gas metabolism, inorganic sulfur oxidation, dissimilatory nitrogen metabolism, and aerobic respiration. **C**: Copy number of the metabolic marker genes encoded in the MAGs. **D**: Copy number of carbohydrate-active enzymes categorized in each MAG, classified as glycoside hydrolases (GHs), glycosyltransferases (GTs), polysaccharide lyases (PLs), carbohydrate esterases (CEs), and auxiliary activities (AAs).

### Core metabolic capacities encoded in the keystone microorganisms

From a community perspective, microbial communities from these three reactors are capable of autotrophic ammonia oxidation (encoding AmoA, RbcL), denitrification (NarG, NirK, NorB, NosZ), sulfur oxidation (FCC, Sqr) as well as trace gas utilization capabilities through high affinity group 1h [NiFe] hydrogenases and carbon monoxide dehydrogenases (CoxL) (**Fig**. 3B, **Tab**. S1). This revealed metabolic flexibility assisted the core functioning of acidic nitritation. A key objective in acidic nitritation is to suppress nitrite oxidizing bacteria (NOB), and our study provides the first metagenomic proof-of-absence of canonical NOB in the reactors, supported by the absence of nitrite oxidoreductase (Nxr) (**Tab**. S2).

*Ca.* Nitrosoglobus kelleri conserves energy via ammonia oxidation to hydroxylamine (NH_2_OH) utilizing ammonia monooxygenase (AmoCAB operon) (**Fig**. 4, **Tab**. S2). The subsequent conversion from NH_2_OH to nitric oxide (NO) was catalysed by the hydroxylamine-ubiquinone redox module (HURM)^59^, consisting of hydroxylamine oxidoreductase HaoAB and the quinone-reducing tetraheme cytochrome cM552 (CycB). Nitrosocyanin has been suggested to catalyze the oxidation of NO to NO_2_ ^-^, which is consistent with that of RJ19 and TAO100^6,60,61^. *Ca.* Nitrosoglobus kelleri encode a NO reductase large subunit (NorB, work in reverse direction) and a putative cytochrome P460, which mediate the production of nitric oxide (N_2_O) from nitric oxide (NO), which was also shown in RJ19 and TAO100^6,62^. An ammonia permease responsible for the uptake of ammonia was detected, congruent with the finding of Picone and colleagues (2021)^6^. Ureolysis produces ammonia and CO_2_ to support chemoautotrophic growth in acid-tolerant AOB. *Ca*. Nitrosoglobus kelleri encodes a complete urease operon (UreABCDEFG), which is similar to TAO100^63^, but lacks the urea transport that was identified in the genome of RJ19^64^.

**Fig. 4.**
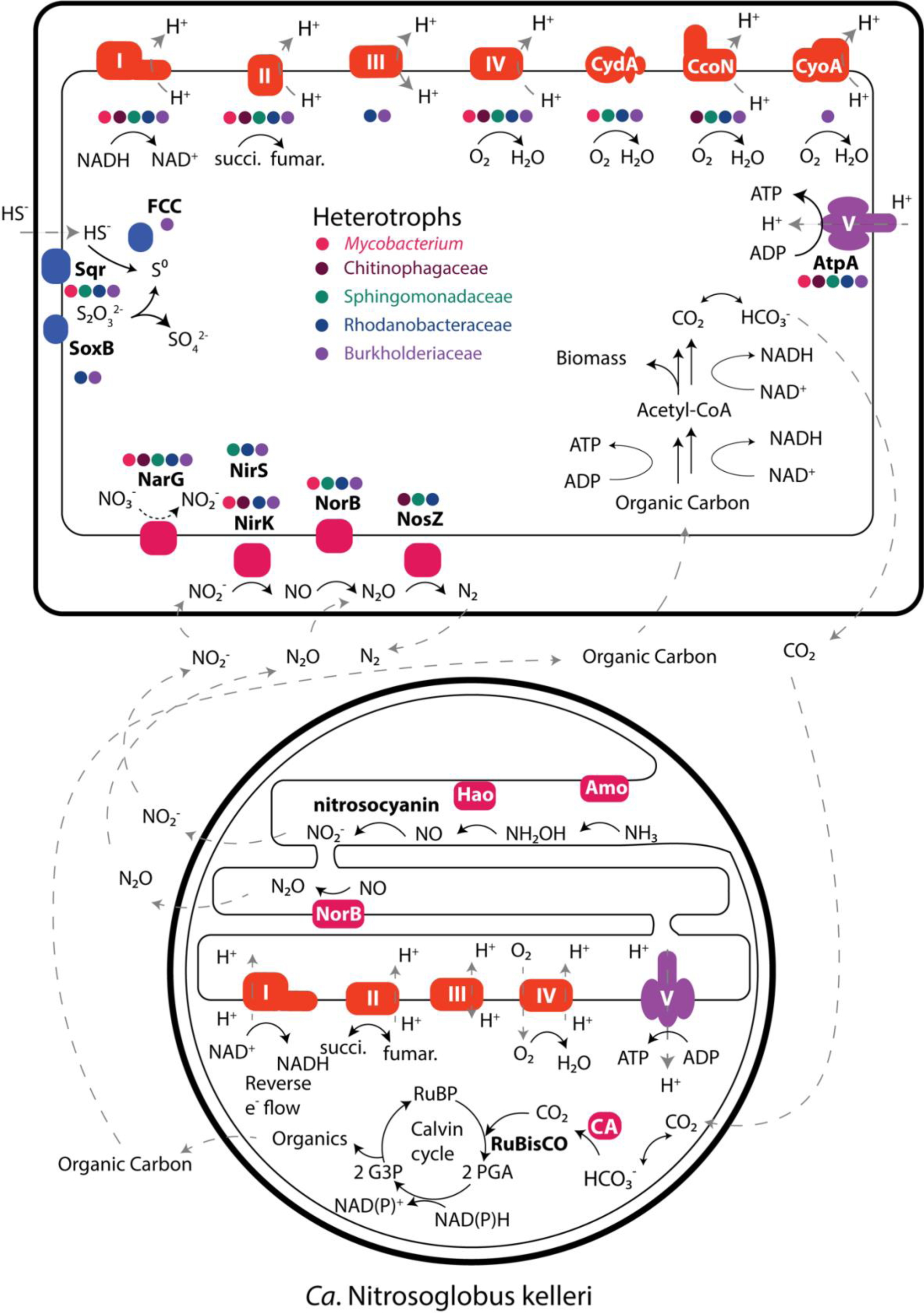
Metabolic interactions between major heterotrophic flanking community members (**top**) and *Ca.* Nitrosoglobus kelleri (**bottom**). The presence of key metabolic genes is summarized for *Mycobacterium* genus, and Chitinophagaceae, Sphingomonadaceae, Rhodanobacteraceae and Burkholderiaceae family. Enzyme complexes are inferred from the presence of the key subunit.

Due to the high redox potential from ammonia oxidation, electrons enter respiratory complex I in *Ca.* Nitrosoglobus kelleri, which works in reverse to generate NADH that fuels carbon fixation^65^, which has been seen in nitrifiers such as *Nitrosococcus oceani*^66^. Different from that of the RJ19 and TAO100, genomes of *Ca.* Nitrosoglobus kelleri contains the complete gene set for succinate dehydrogenase (SDH, complex II), suggesting the presence of complete tricarboxylic acid (TCA) cycle that fuels the respiratory chain.

In contrast to most AOB that encodes form IA RubisCO^67^ for carbon fixation via the complete Calvin–Benson–Bassham (CBB) cycle, *Ca.* Nitrosoglobus kelleri encodes form ID RubisCO, consistent with its proteobacterial origin^68^; except for R2-bin10 that encodes both form ID and IA. This diversification of RubisCO in AOB may reflect selection pressure under acidic nitritation conditions when CO_2_ is potentially limiting. CO_2_ from the environment enters cytoplasm and forms HCO_3_^-^ due to the circumneutral cytoplasmic pH^1^. Carbonic anhydrases function as CO_2_ concentrating mechanisms (CCM)^68^ that converts bicarbonate back to CO_2_ while simultaneously scavenge cytoplasmic protons to mediate pH homeostasis, which was also seen in RJ19 and TAO100^5,6^. Enzymes for carbon storage including phosphoglucomutase, glucose-1-phosphate adenylyltransferase and glycogen synthases were also identified. Genes encoding the biosynthesis of 20 amino acids are complete except for methionine that lacks the enzymes forming cystathionine and homocysteine from either cysteine or aspartate, which was also reported in the genome of RJ19^6^. As for *Mycobacterium* populations, they primarily grow on organic carbon. Surprisingly, R2-bin4 and R3-bin3 encodes high affinity group 1h [NiFe] hydrogenases (Hhy) coupled with form IE RubisCO^69^ suggesting capacity for mixotrophic persistence and growth when organic carbon is limited^70–72^. R2-bin3, R3-bin2, and R3-bin5 further encode actinobacterial aerobic carbon monoxide dehydrogenases (CoxL), further suggest that metabolic flexibility facilitate *Mycobacterium* populations to persist and thrive in the harsh acidic nitritation conditions.

### Metabolic interaction between AOB and heterotrophs

To survive in environments having high acidity and nitrosative stress, reactive nitrogen species (RNS) such as nitrite (NO_2_ ^-^), nitric oxide (NO), and nitrous oxide (N_2_O) generated from ammonium oxidation by *Ca*. Nitrosoglobus kelleri must be consumed. Members of *Mycobacterium* and Rhodanobacteraceae, Burkholderiaceae, Chitinophagaceae, and Sphingomonadaceae encode complete or truncated denitrification machinery to consume reactive nitrogen species to mitigate nitrosative stress (**Fig**. 4).

In terms of the carbon cycle, since the synthetic wastewater in R2 contains no organic carbon, it is clear that autotrophic *Ca.* Nitrosoglobus kelleri assimilates carbon dioxide into biomass and synthesizes amino acids to support heterotrophic growth in *Mycobacterium*, Rhodanobacteraceae and Burkholderiaceae. Feed of R2 is free from organic carbon, carbon dioxide produced during aerobic respiration from heterotrophs in turn, is the sole carbon source for *Ca.* Nitrosoglobus kelleri in this regard (**Fig**. 4).

Alternatively, since municipal wastewater typically contains high levels of bioavailable organic carbon (e.g., influent of R1 contains soluble COD at around 200 mg/L^10^), it supports heterotrophic growth as well. The complexity of organic input in the wastewater is reflected by the large diversity of CAZymes that are responsible for the degradation of diverse organic compounds that exist in wastewater, which also suggest the possibility of metabolic cross feeding of organic compounds in these microbial communities (**Fig**. 3D). Interestingly, the genomes of *Ca.* Nitrosoglobus kelleri encodes 27-28 GT, 12-15 GHs, 7-8 CEs, 4 AAs and 2-3 CBMs (**Tab**. S2), which infer capacity cellular maintenance e.g., assimilatory recycling of organic materials.

### Resilience in *Ca.* Nitrosoglobus kelleri and *Mycobacterium*

Acidic nitrification is a unique ecosystem, where the protonation of ammonia leads to decreased substrate availability and formation of toxic nitrogenous compounds creates cellular nitrosative stress^5^, in addition to the need to maintain cytoplasmic homeostasis caused by protons intrusion at low pH. We compared the acid-tolerant AOB genomes i.e., *Ca.* “*Nitrosoglobus kelleri*”, RJ19 and TAO100 against four neutrophilic representative species of *Nitrosococcaceae* available on NCBI or GTDB (R207_2, accessed on Oct 19, 2022). In particular, a H^+^/Na^+^ antiporter (NhaA) homologous to that of *Acidithiobacillus* (an obligate acidophilic chemolithotrophic genus), which uptake Na^+^ in exchange of cellular H^+^ from the cytoplasm is exclusively present in these three genomes (**Tab**. S3). Multiple cation transporters were also found, which creates Donnan potential across the membrane resulting in a chemiosmotic gradient that inhibits proton influx (**Tab**. S3)^7^. We also found major facilitator superfamily (MFS) proteins are present and suggested to mediate the transport of drugs, metabolites, and ions as well as mitigating oxidative stress^73,74^. Glutathione (GSH) production capacity was also encoded exclusively in these three genomes, a ubiquitous tripeptide thiol, is a vital antioxidant to provide cellular defence against oxidative stresses^75,76^. Multiple genes were identified and confer roles in cellular envelope biosynthesis, which confer capacity for cells to build a stronger cellular envelope to defend against the destructive effects posed by FNA on cellular envelope materials e.g., lipopolysaccharides, peptidoglycan, and teichoic acid polymers and extracellular polymeric substances (EPS)^77,78^ (**Tab**. S3).

Similarly, *Mycobacterium* have recognized cell wall^79^ and cytoplasmic membrane^80^ highly impermeable to proton intrusion, which facilitate their persistence during acidic nitritation. We further identified F_0_F_1_-type ATP synthases (**Fig**. 4) in *Mycobacterium* MAGs that are crucial for growth and pH regulation^81,82^ (**Tab**. S4). A membrane associated oxidoreductase complex (MRC) that is crucial in coordinating radical detoxification systems enabling *Mycobacterium tuberculosis (Mtb)* to survive the oxidative stress imposed by host phagocytes^83^ was shown to be well-conserved in all *Mycobacterium* MAGs in this study (Tab. S4). In addition, multiple potassium transport systems, including potassium-transporting ATPase, Trk system and K^+^/H^+^ antiporters^84–86^, couple with multiple sodium/proton antiporters^87,88^ contribute to the regulation of pH homeostasis through the establishment of internally-positive membrane potential, which is also seen in *Ca*. Nitrosoglobus kelleri (**Tab**. S4)^7,88,89^. These mechanisms work in symphony with proton consumption reactions mediated by carbonic anhydrases^2,6,86^ and glutamate decarboxylases^88,90^, as well as DNA repair systems to regulate intracellular pH and repair DNA damage caused by acidic and nitrosative stress (**Tab**. S4)^7,89,91^.

In addition to trace gas oxidation capability mentioned above, inorganic sulfur oxidation, including sulfide oxidation to elemental sulfur mediated by sulfide-quinone oxidoreductase (Sqr) are identified in Sphingomonadaceae, Rhodanobacteraceae, Burkholderiaceae, and *Mycobacterium*; flavocytochrome c flavoprotein (FCC) was identified in Burkholderiaceae; while thiosulfate oxidation mediated by the Sox complex that have been identified in Rhodanobacteraceae and Burkholderiaceae. Hydrogen sulfide (H_2_S) is a common contributor of sewage odor^92^, it’s possible that these heterotrophs use inorganic sulfur to conserve energy. Therefore, it is apparent metabolic flexibility, such as using alternative energy sources, contribute to the energetically costly pH homeostasis and oxidative stress mitigation processes.

## Conclusions

Our analysis has focused on depicting an ecological model in which the intrinsic nitrification activity of *Ca*. Nitrosoglobus kelleri creates an environment with high acidity and oxidative stress, while *Mycobacterium* populations are able to survive in this harsh environment while detoxifying reactive nitrogen species. To date, no other microorganism was reported to be able to mediate such reciprocal relationships and enabled the functioning of acidic nitritation^17^. These insights contribute to the development of theoretical frameworks in microbial ecology. It is also the foundation of an innovative wastewater treatment process that is promising in reducing the environmental impact. In light of this work, obtaining pure cultures of these organisms, followed by genetic manipulation to further validate the findings from this study is warranted. In turn, this will provide new understandings of how life persists and grows in extreme environments.

### Etymology

*Candidatus* (*Ca*.) Nitrosoglobus kelleri sp. nov. We name this acid-tolerant AOB *Ca*. Nitrosoglobus kelleri after Professor Jurg Keller at the University of Queensland in respect to his distinguished contributions to our progress of a more sustainable water industry.

## Ethics declaration

These authors declare no competing interests.

## Acknowledgement

This work was supported by the Advance Queensland Industry Research Fellowship (RM2019002600) and the Early Career Postdoctoral Fellowship (ECPF23-8566329039) at the Faculty of Medicine, Nursing and Health Sciences of Monash University awarded to G. Ni. C. Greening is supported by an Australian Research Council (ARC) Discovery Project (DP210101595). This work was also supported by National Health & Medical Research Council (NHMRC) grant APP1178715 to C. Greening. J. Guo and M. Zheng acknowledge the support of an ARC Discovery Project (DP230101340). M. Zheng also acknowledges the support of an ARC Industry Fellowship (IE230100245).

---

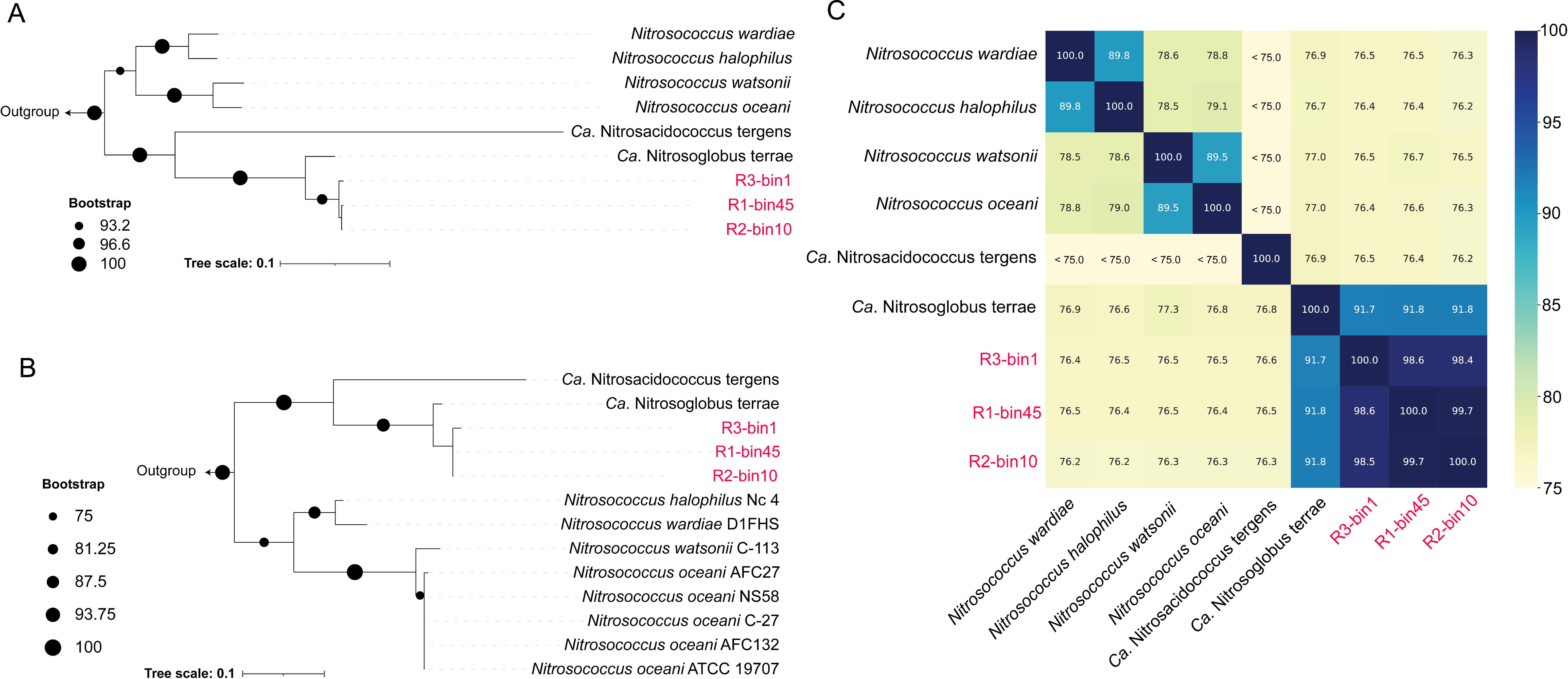

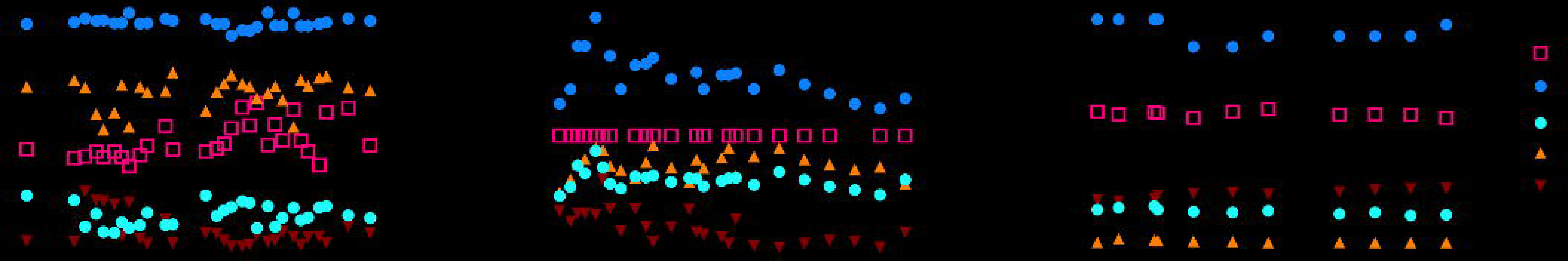

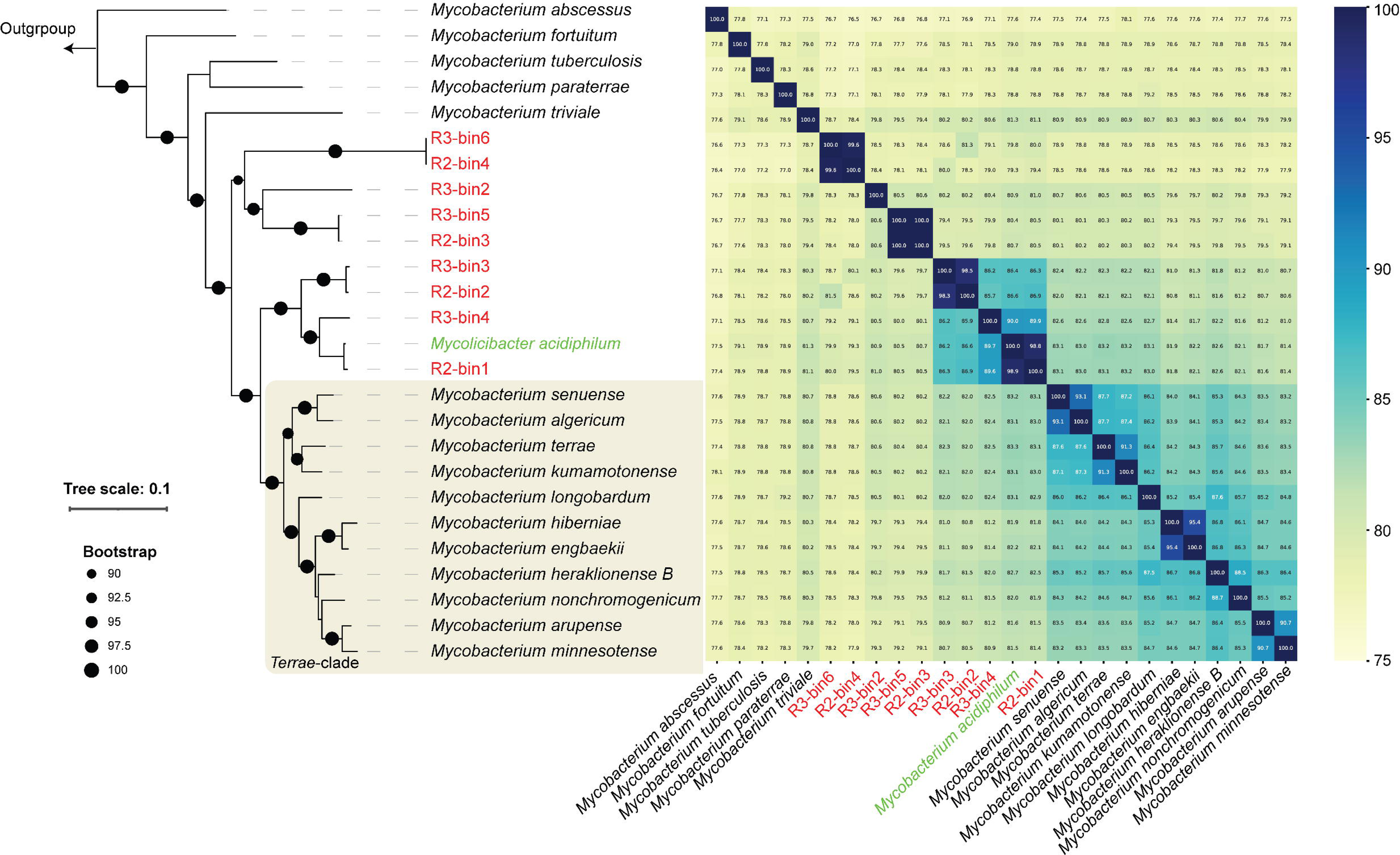

## Notes

### Competing Interest Statement

The authors have declared no competing interest.

